# Cross-kingdom analysis of microbial communities in Cystic Fibrosis and Bronchiectasis

**DOI:** 10.1101/2022.01.11.475678

**Authors:** L. Cuthbertson, J. Ish-Horowicz, I.C. Felton, P. James, E. Turek, M.J. Cox, M.R. Loebinger, N.J. Simmonds, S. Filippi, M.F. Moffatt, W.O.C. Cookson

**Author notes:** Authors contributed equally to this manuscript (L. Cuthbertson, J. Ish-Horowicz). Joint senior authors (M.F. Moffatt, W.O.C. Cookson). **Corresponding author:** L. Cuthbertson, National Heart and Lung Institute, Imperial College London, London SW3 6LY, UK.

## Abstract

**Background:** Cystic fibrosis (CF) and non-CF bronchiectasis (BX) are characterised by severe chronic infections. Fungal and bacterial components of infection are both recognized. Little however is known about how fungal and bacterial organisms interact and whether these interactions impact on disease outcomes.

**Methods:** Quantitative PCR and next-generation sequencing of ITS2 and 16S rRNA gene was carried out on 107 patients with CF or BX with clinically defined fungal infection status for all patients. The relationship between fungal and bacterial community composition was extensively explored using: random forest modelling, correlation network analysis, multi-omics factor analysis, and sample-wise clustering, to understand associations both within and between the microbial communities and their relationship to respiratory disease.

**Results:** Random forest modelling demonstrated distinct fungal and bacterial communities within CF and BX patients. The inclusion of both kingdoms in the models did not improve discrimination between the two diseases. Within the CF patients, bacterial community composition was independent of clinical fungal disease status. Bacterial and fungal communities did not relate to the presence of CF pulmonary exacerbations (CFPE). Correlation network analysis found intra-kingdom interactions were predominant in the data. Multi-omics factor analysis (MOFA) revealed latent factors corresponding to single kingdoms. Thus, in the bacterial community we identified two distinct clusters characterised by the presence or absence of *Pseudomonas*-domination. This was independent of fungal community which was characterised by a second set of independent clusters dominated by *Saccharomycetes*.

**Conclusions:** In this study we were unable to detect clear evidence of clinically significant inter-kingdom interactions between the bacterial and fungal communities. While further work is required to fully understand microbial interaction within the lung, our data suggests that interkingdom interactions may not be the primary driver of patient outcomes, particularly in the context of fungal infection.

## Introduction

Chronic respiratory infections are the leading causes of morbidity and mortality for the chronic suppurative lung diseases (CSLD) cystic fibrosis (CF) and non-CF bronchiectasis (BX) (1, 2). Bacterial infections are a major pathophysiological factor in disease progression in these patients (3, 4). The impact of fungal infections is increasingly being recognised and fungal infections have been associated with higher disease burdens, increased exacerbation rates, and accelerated clinical decline (5, 6).

The development of robust microbial sequencing protocols has revealed complex fungal communities within the lungs of patients with CF and BX, exhibiting a range of clinical disease manifestations (7, 8). The recognition of these fungal communities has led to the investigation of bacterial and fungal interactions and their roles in disease progression. Insights into these complex associations and interactions are helped through evolving statistical methodology (8).

Machine learning techniques have proven to be effective for host phenotype prediction from microbiome profiles with random forests exhibiting the strongest predictive performance in host-trait prediction tasks (9). This superior performance is driven in part by their ability to model non-linear interactions between variables. This property is especially useful in microbiome studies, in which the covariates represent a dynamic and interacting ecological system of microbes. Random forests are therefore often preferred over more interpretable linear models, with a recent systematic review finding that random forest was the most popular machine learning model for differential abundance testing in microbiome studies (10).

In this study we aimed to explore fungal and bacterial interactions within the lung of patients with chronic respiratory diseases (CF and BX) and understand their relationship to underlying disease, fungal disease diagnosis and CF pulmonary exacerbations (CFPE). Using random forest analysis we explored the predictive power of bacterial and fungal community composition and the presence of CF (n=83) or BX (n=24). This pipeline was subsequently applied to two fungal disease sub-groups within the CF disease group (see Table 1), defined by the presence (n=20) or absence (n=39) of clinical diagnoses of fungal bronchitis (FB). Due to the small number of patients with BX (n=24), we confined this sub-group analysis to CF patients only. Motivated by recent work by Soret *et al*. (8), we also investigated CFPEs.

**Table 1.** Patient demographics. Data for all samples included in this manuscript. Only samples with paired fungal and bacterial sequencing that passed all quality control steps (see Supplementary Material) were taken forward for analysis. Differences in continuous variables were calculated using a one-way t-test, while a chi-square test was used for categorical variables.

|  | **BX** (n = 24) | **CF** (n = 83) | *P* |
| --- | --- | --- | --- |
| Age/yrs (mean [SD]) | 59.79 (11.06) | 32.00 (11.90) | <0.001 |
| Male (%) | 10 (41.7) | 47 (56.6) | 0.288 |
| BMI (mean [SD]) | 26.51 (8.73) | 22.51 (6.50) | 0.029 |
| FEV<sup>1</sup> % predicted (mean [SD]) | 58.24 (25.23) | 48.81 (17.79) | 0.047 |
| Disease Group (%) |  |  | <0.001 |
| *ABPA* | 8 (33.3) | 14 (16.9) |  |
| *CNPA* | 5 (20.8) | 0 (0.0) |  |
| *FB* | 0 (0.0) | 20 (24.1) |  |
| *NAFD* | 8 (33.3) | 39 (47.0) |  |
| *NTM* | 3 (12.5) | 10 (12.0) |  |
| Exacerbation* (%) | 1 (4.2) | 36 (43.4) | 0.001 |

CF: Cystic fibrosis, BX: Non-CF bronchiectasis, BMI: Body mass index, FEV<sup>1</sup>:
Forced expiratory volume in 1 second, ABPA: Allergic bronchopulmonary
aspergillosis, CNPA: Chronic necrotising pulmonary aspergillosis, FB: Fungal
bronchitis, NAFD: No active fungal disease, NTM: Non-tuberculosis mycobacteria.
\* Patients were defined as exacerbating by a change in clinical status from
baseline that resulted in a new pulmonary antibiotic treatment.

The random forest analyses were complemented by unsupervised approaches investigating both intra- and inter-kingdom associations between taxa (correlation network analysis and multi-omics factor analysis (11)) and samples (Dirichlet multinomial mixtures (12)). The focus of both sets of the analyses was to find evidence of interactions between the two kingdoms.

## Methods

### Study Design

A prospective, cross-sectional study of spontaneously expectorated sputum from adults with CF and BX was carried out at the Royal Brompton NHS trust between April 2013 and July 2014 (7). Ethical approval was obtained from the Royal Brompton and Harefield Hospital Biomedical Research Unit Ethics Committee (Advanced Lung Disease Biobank study number: 10/H0504/9). Written informed consent was obtained from all study participants prior to sample collection. This study was conducted in accordance with the International Conference for Harmonisation of Good Clinical Practice and the guiding principles of the Declaration of Helsinki and the Research Governance Framework for Health and Social Care.

Patient details, including treatments, were collected and subjects were partitioned into defined clinical subgroups according to the diagnostic criteria defined in Cuthbertson *et al*. 2020 (7) (Table 1). Participants within the study cohort were classified into four clinically defined fungal disease groups (see Table 1): (i) fungal bronchitis (FB), (ii) Allergic bronchopulmonary aspergillosis (ABPA), (iii) chronic necrotising pulmonary aspergillosis (CNPA, BX only); and (iv) non-tuberculous mycobacteria (NTM).

Sputum samples were collected and processed as previously described, with half the sample sent for routine clinical microbiological culture and the other half stored at −80°C for DNA analysis (7). DNA extraction was performed using the DNA fast spin kit for soil (MPBio, California, USA) according to the manufacturer’s instructions. Extraction controls were blinded and processed along with patient samples (7).

### Quantitative PCR

Bacterial biomass was quantified by SYBR green quantitative PCR (qPCR) (13). Fungal biomass was estimated using a modified Taqman based qPCR assay as previously described (7). All qPCR reactions were performed in triplicate.

### DNA sequencing

16S rRNA gene and ITS2 sequencing were performed on the Illumina MiSeq platform using dual barcode fusion primers. Bacterial sequencing was performed on the V4 region of the 16S rRNA gene as previously described (13). ITS2 sequencing was performed using the primers, ITS2F (5’-CAR CAA YGG ATC TCT TGG-3’) and ITS2R (5’-GAT ATG CTT AAG TTC AGC GGG T-3’) with ligated adaptors (7). Extraction controls, PCR negative and mock communities were included on all sequencing runs.

Sequence processing of 16S rRNA gene and ITS2 data was carried out using QIIME 1.9 as described previously in Cuthbertson *et al*. 2017 (13) and Cuthbertson *et al*. 2020 (7) respectively.

All sequences were submitted to the European nucleotide database. Bacterial data can be accessed under project number PRJEB33064 with the fungal data accessible under project number PRJEB33434.

### Statistical analysis

Statistical analysis was carried out in R version 3.5.1. Data was analysed in phyloseq version 1.24.2 (14). Decontamination of the data was carried out using decontam version 1.1.2 (15); full details are available in Supplementary Material. Differences between categorical variables were calculated using Wilcoxon rank sum test and Kruskal-Wallis. Pearson correlation was used for tests between continuous variables. Differences in microbial community composition were tested with PERMANOVA using the Adonis function from vegan version 2.5-6 [23]. Random forests models were fitted using the caret (version 6.0) and ranger (version 0.12) packages 4. The DirichletMultinomial package (version 1.36) was used for sample-wise clustering (16). MOFA analysis used the MOFA2 package (version 1.7) (11). *P*-values were adjusted for multiple corrections using false discovery rate (FDR) throughout.

### Random forest-based two-sample testing

Random forest (RF) binary classifiers were used to investigate associations between microbial community composition and the three groups described in Table 2. We agglomerated the OTU counts to each rank in Class, Order, Family, Genus and Species. For each rank we trained three models with the following covariates:

**Table 2.** Patient groups investigated using random forest. These groups were chosen to maximise sample sizes and power of the analyses. The Group phenotype was limited to the FB (n = 20) and NAFD (n = 39) groups within the CF population. BX samples were removed from the group analysis due to the low numbers.

| Group | Description | Classes | n |
| --- | --- | --- | --- |
| Disease | Whether a patient has CF or BX | CF or BX | 83 CF, 24 BX (107 total) |
| Fungal disease (CF only) | Whether a CF patient has been diagnosed with FB or NAFD | FB or NAFD | 20 FB, 39 NAFD (59 total) |
| CFPE (CF only) | Whether a CF patient was experiencing pulmonary exacerbation when the sample was collected (clinician defined). | Yes or No | 36 Yes, 47 No (83 total) |

CF: cystic fibrosis, BX: non-cystic fibrosis bronchiectasis, FB: fungal bronchitis,
NAFD: no active fungal disease.

1. the agglomerated counts of the bacterial OTUs;
2. the agglomerated counts of the fungal OTUs; or
3. the agglomerated counts of both the bacterial and fungal OTUs.

The five different agglomeration ranks were used in order to investigate the differences between groups at different levels of the taxonomic hierarchy. For all models, any agglomerated taxa present in fewer than 20% of samples were removed. Removal of rare samples has been shown to improve stability and reproducibility of random forest analyses while still retaining discriminative taxa and leaving predictive performance unchanged (17).

The predictive performance of each RF model was estimated using 5-fold nested cross-validation with 5 folds in the inner loop. The inner loop performed model selection using a random hyperparameter search of ten combinations of mtry and splitrule. Both inner and outer folds were sampled in a stratified manner meaning that the class proportions in each fold reflected the proportions of the entire dataset. The predictive performance of the models was evaluated using area under receiver-operating characteristic (ROC) and precision-recall (PR) curves. Ninety-five % confidence intervals on the cross-validated area under the curves (AUCs) were computed using the precrec package (18). The statistical significance of AUC values were computed using a label permutation test, where observed AUCs were compared to AUCs from random forest models trained on permuted labels. The dependence of the random forest results to the number of outer-loop folds was investigated by repeating all analyses using 10 outer folds (Supplementary Table S1).

Similar results for this dataset showing the robustness of the predictive metrics (area under ROC and PR curves) and the increased stability of variable importance scores under different transformations are included in the Supplementary Material (Table S1 and Figure S1-5). For the results in the main text, taxa reads were converted to relative abundances by dividing by the total per-sample reads. The random forest AUC and variable importance results were robust under four additional transformations (identity, centre log-ratio (CLR), log(x+1), and division by the sum of dataset reads). Please see Supplementary Table S1 for AUC results for the random forest models under the different transformations.

### Random forest variable importance and differential abundance analysis

One of the primary benefits of random forest modelling is the ability to perform a variable importance analysis. Here, we use both of the two most popular variable importance measures for random forest - Mean Decrease Gini and Mean Decrease Accuracy. In addition, we include the de-biased Mean Decrease Gini scores proposed by Nembrini *et al*. (19).

Each random forest was grown using 1,000 trees to achieve stability in the variable importance scores. The statistical significance of MDA and de-biased MDG scores were assessed using *P*-values calculated using the permutation method of Altmann *et al*. with 1,000 permutations (20). Random forest variable importance scores are unsigned (do not give an effect direction), although an approximate effect direction can be obtained using partial dependence plots. Such plots visualise the marginal effect of a variable on the predictions of a model and are used here to gain further insight into the dependence of model prediction on the relative abundance of different taxa (21).

### Correlation analysis, MOFA and sample-wise clustering

We analysed the correlation structure amongst the taxa included in the random forest analysis. Samples were centred log-ratio (CLR) transformed prior to calculating correlations to account for the compositional nature of 16S rRNA gene and ITS2 sequencing data. The CLR transformation was applied separately to the 16S rRNA gene and ITS2 samples, as has been previously applied to find accurate cross-omic interactions (22). This pre-transformation requires that the resulting correlations are interpreted as the log-ratios abundance relative to the sample geometric mean rather than in absolute terms. MOFA was run using the same pre-processing as Haak *et al*. (23) (agglomeration to Genus and transformation using CLR prior to analysis). Sample-wise clustering was performed using Dirichlet Multinomial mixtures (12) on the un-transformed reads of each kingdom separately. The number of clusters was selected by comparing the Akaike and Bayesian information criteria and the Laplace approximation of the model evidence for 1 to 15 clusters.

## Results

### Description of data

After decontamination and removal of samples with less than 2,000 reads (n = 2), 107 samples were retained for downstream analyses (for demographics see Table 1). For the ecological analyses, bacterial reads were rarefied to 2,357 while fungal reads were rarefied to 2,542. All other analyses used un-rarefied reads.

## Ecological analysis

### Differences between diseases

We performed an ecological analysis on the rarefied data (full details are available in Supplementary Material). In short, Wilcoxon rank sum tests revealed both bacterial and fungal diversity were significantly higher in patients with BX than CF (*P* < 0.001). Similarly, bacterial biomass was significantly higher in the BX group (W = 1,100, effect size = 0.241, *P* = 0.018) but no significant difference in fungal biomass was observed. PERMANOVA revealed significant but small differences in community composition between CF and BX (bacterial, R^2^ = 0.066, *P* <0.001; fungal R^2^ = 0.028, *P* = 0.004).

### Differences between fungal disease groups in cystic fibrosis

Within the CF group, we compared the two largest fungal disease groups, FB (n=20) and NAFD (n=39). We observed significant differences in fungal alpha diversity between the NAFD and FB groups (Wilcoxon rank sum test, *P* < 0.05). There were no significant differences, however, in bacterial biomass or diversity.

### CFPE

There was no significant difference in bacterial or fungal, biomass or alpha diversity measures (Wilcoxon rank sum test, *P* > 0.1) between CFPE subjects and those that were stable.

## Random forest analysis

### Discriminative power of bacterial and fungal communities

We further investigated differences in fungal and bacterial community composition between groups of patients using random forest modelling (Figure 1). For each set of group labels, we trained a random forest binary classifier with covariates being OTU reads agglomerated to Class, Order, Family, Genus, or Species level. Differences between groups were quantified using the area under ROC or PR curve (AU-ROC and AU-PRC). The null hypothesis (no difference between the groups) implies an AU-ROC=0.5 and an AU-PR equal to the proportion of the positive class. The baseline AU-PRC therefore varies between the different sets of group definitions.

**Figure 1:**
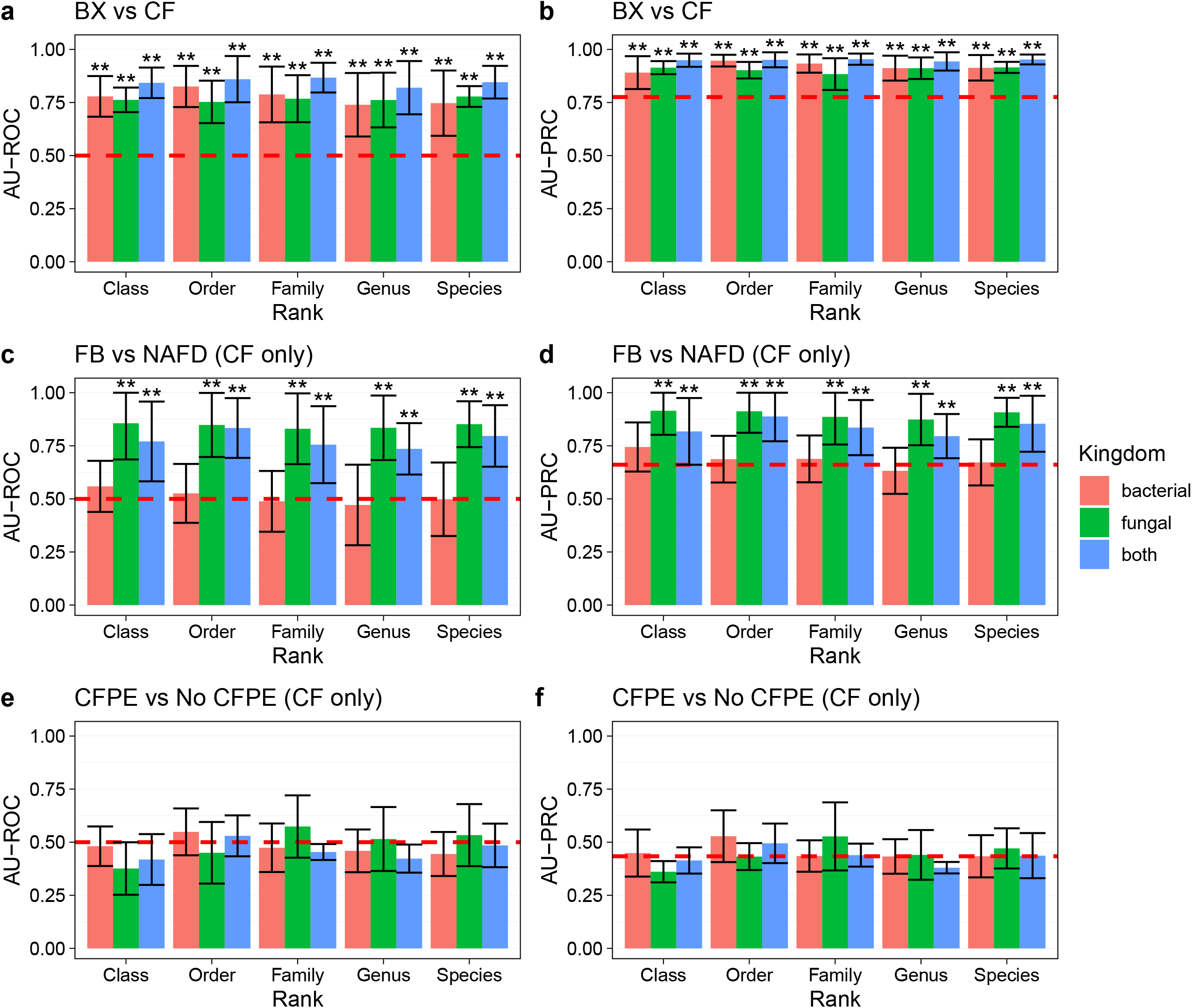
Discriminative power (quantified using area under ROC and PR curve) of random forest models predicting (a,b): disease status; (c,d): fungal disease status (CF only); and (e,f) and CFPE (CF only). The expected values for a random classifier (indicating no difference between groups) is denoted by a black dotted line. For AU-ROC (plots a,c,e) this is 0.5, while for AU-PRC (plots b,d,f) while for AU-PRC the value is the proportion of samples in the positive class. Error bars are 95% confidence intervals. *: *P* < 0.10, **: *P* < 0.05 for 100 replicates of a label permutation test. P-values adjusted using false discovery rate.

Statistically significant differences between the CF and BX groups were evident at every taxonomic rank. These differences were detected using both the AU-ROC (Figure 1a) or AU-PRC (Figure 1b) metrics. We also found that the differences between CF and BX did not differ significantly between the two kingdoms (label permutation test, *P* > 0.10), meaning that both kingdoms have equally distinct communities between CF and BX. In addition, including both communities in a random forest does not increase the predictive power (label permutation test, *P* > 0.10).

Figures 1c and 1d show that fungal disease group within the CF patients is independent of bacterial community composition. As expected, fungal community composition is a good predictor of fungal disease status within the CF group. Adding the bacteria to the random forest model decreased predictive power at all ranks. The bacterial covariates represent additional noise in the context of fungal disease group however, the models still have better than-random performance due to the inclusion of the fungal taxa (label permutation test, *P* < 0.05).

Finally, CFPE was found to be independent of both bacterial and fungal community composition (Figures 1e and 1f), none of the random forest models had predictive power significantly better than random (label permutation test, *P* > 0.10).

### Differential abundance analysis

Random forest models that detected a significant difference between their two classes were then analysed using three variable importance measures: mean decrease accuracy (MDA); mean decrease Gini (MDG); and de-biased mean decrease Gini (corrected MDG, (19)). Statistical significance can only be assessed for the MDA and de-biased MDG scores. Only Genus-level agglomeration was included as it is the highest reliable taxonomic resolution for both 16S rRNA gene and ITS2 sequencing (24). The four random forest models that detected a difference between their respective groups were:

1. distinguishing CF/BX using bacterial and fungal genera;
2. distinguishing CF/BX using bacterial genera;
3. distinguishing CF/BX using fungal genera; and
4. distinguishing FB/NAFD using fungal genera (CF group only).

The most highly-ranked taxa for these four models are shown in Figure 2(a-d). For Model 1, *Penicillium* (direction BX) is the most highly-ranked genus according to all three of the variable importance methods. *Pseudomonas* (direction CF), *Malassezia* and Neisseria (direction BX) are also highly-ranked. All four of these associations are significant (p<0.05) when using the de-biased MDG scores, *but not for MDA scores*.

**Figure 2:**
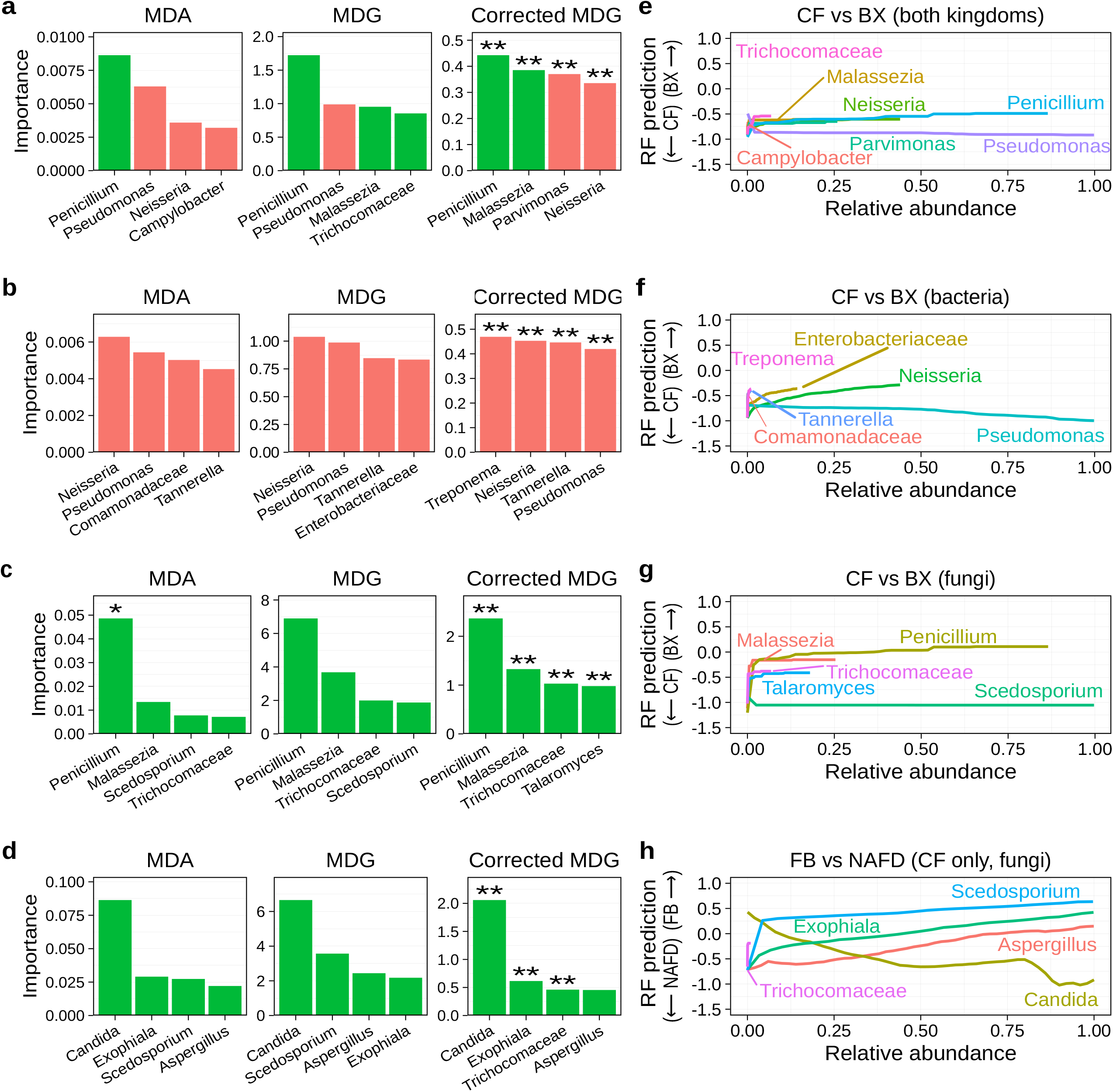
Each row shows the four top-ranked taxa and partial dependence for a random forest model distinguishing (a,b): BX/CF from bacterial and fungal genera; (c,d): Bx/CF from bacterial genera; (e,f): Bx/CF from bacterial genera; and (g,h): FB/NAFD from fungal genera (CF group only). *: *P* < 0.10, **: *P* < 0.05 for 100 replicates of a label permutation test. *P*-values adjusted using false discovery rate.

Importance rankings can be augmented by partial dependence plots (Figure 2 b,d,f,h) which visualise the marginal effect of a single variable on the prediction of the model. These provide effect directions for the unsigned variable importance scores as well as insight into the type of dependence. For example, the majority of the effect of the important taxa for Model 1 (Figure 2b) occurs when the relative abundance increases from zero to non-zero relative abundance. This is the case for Pseudomonas, where the likelihood of a BX prediction quickly decreases as its relative abundance increases, before the rate of decrease slows.

Model 2 ranks Treponema, Neisseria (direction BX), *Pseudomonas* and Tanerella (direction CF) highly (Figure 2c). Overlap is observed with Model 1 as the two models share bacterial covariates. The partial dependence (Figure 2d) shows that increasing *Pseudomonas* relative abundance does not increase the likelihood of CF until it increases beyond 50%.

Model 3 (Figure 2e), clearly indicates that *Penicillium* is the most important taxa and is significantly associated with an increased probability of BX in this cohort. Once again, there is overlap with Model 1 due to shared fungal covariates. Compared to Model 1, however, this fungal-only model places higher importance on *Penicillium*, which can be seen from the much steeper increase in the partial dependence (Figure 2f) at small relative abundance.

Model 4 (Figure 2g) ranks *Candida* (direction NAFD) as the most important taxa, followed by Exophiala, *Aspergillus* and *Scedosporium* (direction FB). *Candida* is an opportunistic pathogen, but these partial dependence plots (Figure 1h) show a lower likelihood of FB prediction from the model as a sample becomes increasingly *Candida*-dominated.

### Correlation analysis and clustering

Correlation network analysis is a useful tool to explore microbial associations (8, 25, 26). Strong correlations are commonly observed in microbiome studies and correlations can be positive; associated with microbes inhabiting common ecological niches, or negative; indicating competition. In this dataset we observe clear structure at the genus level (Figure 3a), with blocks of positively correlated bacterial (*Steptococcus, Veillonella, Rothia, Selemonas, Prevotella, Fusobacterium, Atopobium, Neisseria, Haemophilus* and *others*) and fungal genera (*Serratia, Talaromyces, Filobasildea, Fusarium* and *Trichosporon*). The genera in these blocks come from a single kingdom and so do not indicate prominent cross-kingdom dependencies in the community structure. In addition, there are no significant correlations between members of these two blocks of taxa (*P* > 0.01), suggesting that the two blocks are largely independent of one another. This may be because they occupy separate niches in the respiratory tract or due to sampling bias.

**Figure 3:**
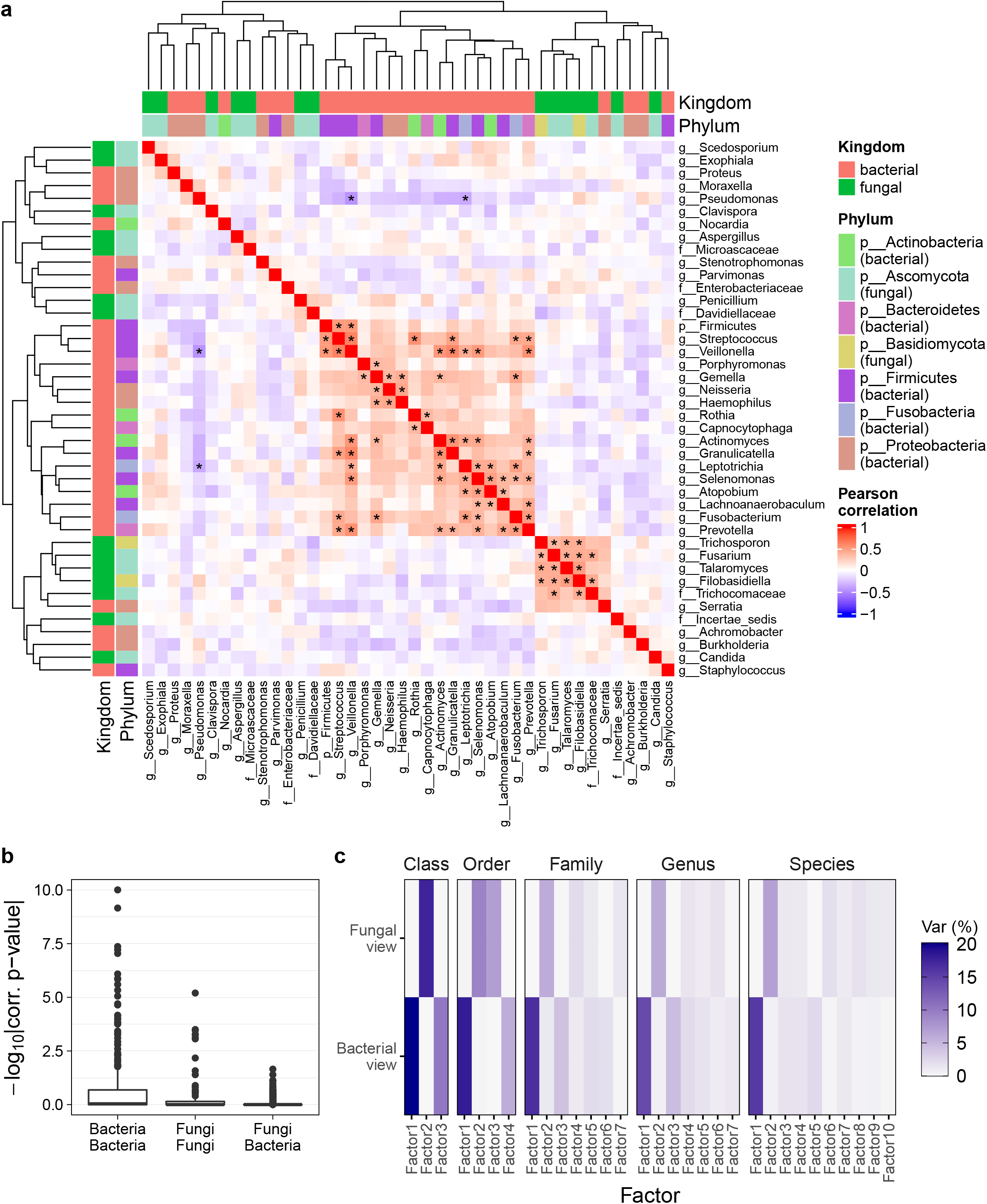
(a) Pearson correlation analysis shows that taxa form correlated clusters with members of the same kingdom. Bacterial and fungal abundances are agglomerated to Genus level and transformed (separately) using centred log-ratio prior to calculating correlations. Genera accounting for more than 0.1% of reads in their respective kingdom are shown. * indicates pairwise correlations for which *P* < 0.01. (b): The *P*-values for the correlations in panel (a) are smaller for correlations within each kingdom than between the two kingdoms (c): Multiomics factor analysis (MOFA) identifies latent factors that describe the variance in the composition of both kingdoms. However, these latent factors each describe variance in a single kingdom.

The relative importance of intra- and inter-kingdom correlations was further explored by considering the *P*-values from the pairwise correlations (Figure 3b), which indicated that significant correlations occur within each kingdom more than between. Figure 3b also shows that, while the most extreme correlations are between pairs of fungal genera, the correlation patterns within the bacterial genera are overall more significant.

A dedicated multi-omics integration approach was used to further investigate the underlying drivers of cross kingdom community structure by applying Multi-omics factor analysis (MOFA, (11)). MOFA is an unsupervised method that finds latent factors that explain the variance across different “views” of the same samples: in this analysis, bacterial and fungal abundances only identify latent factors that explain variance in a single kingdom (Figure 3c). This is true at five different taxonomic levels of agglomeration (Class, Order, Family, Genus, Species). This provides further evidence on the lack of detectable cross-kingdom dependencies in this dataset.

Community structure within the microbial kingdoms across samples was further analysed with Dirichlet mixture components, grouping samples into distinct clusters with similar composition (12). This unsupervised approach provides insight into community-level structure across samples, which may or may not correspond to the pre-defined clinical labels.

All 107 samples were clustered using Dirichlet Multinomial Mixture models using raw count values agglomerated to one of the taxonomic ranks. This was performed separately on the two kingdoms and resulted in two sets of cluster labels for each agglomeration rank. Using information-theoretic goodness of fit measures (Figure S6), two distinct bacterial clusters were found at Genus level and two fungal clusters were found at Class level. Both the bacterial (Figure 4a, top) and fungal (Figure 4a, bottom) clusters are separable in Bray-Curtis principal coordinate space. The clusters for bacterial genera are defined by *Pseudomonas* domination (Figure 4d) while the fungal class clusters are defined by *Saccharomycetes* domination (Figure 4e).

**Figure 4:**
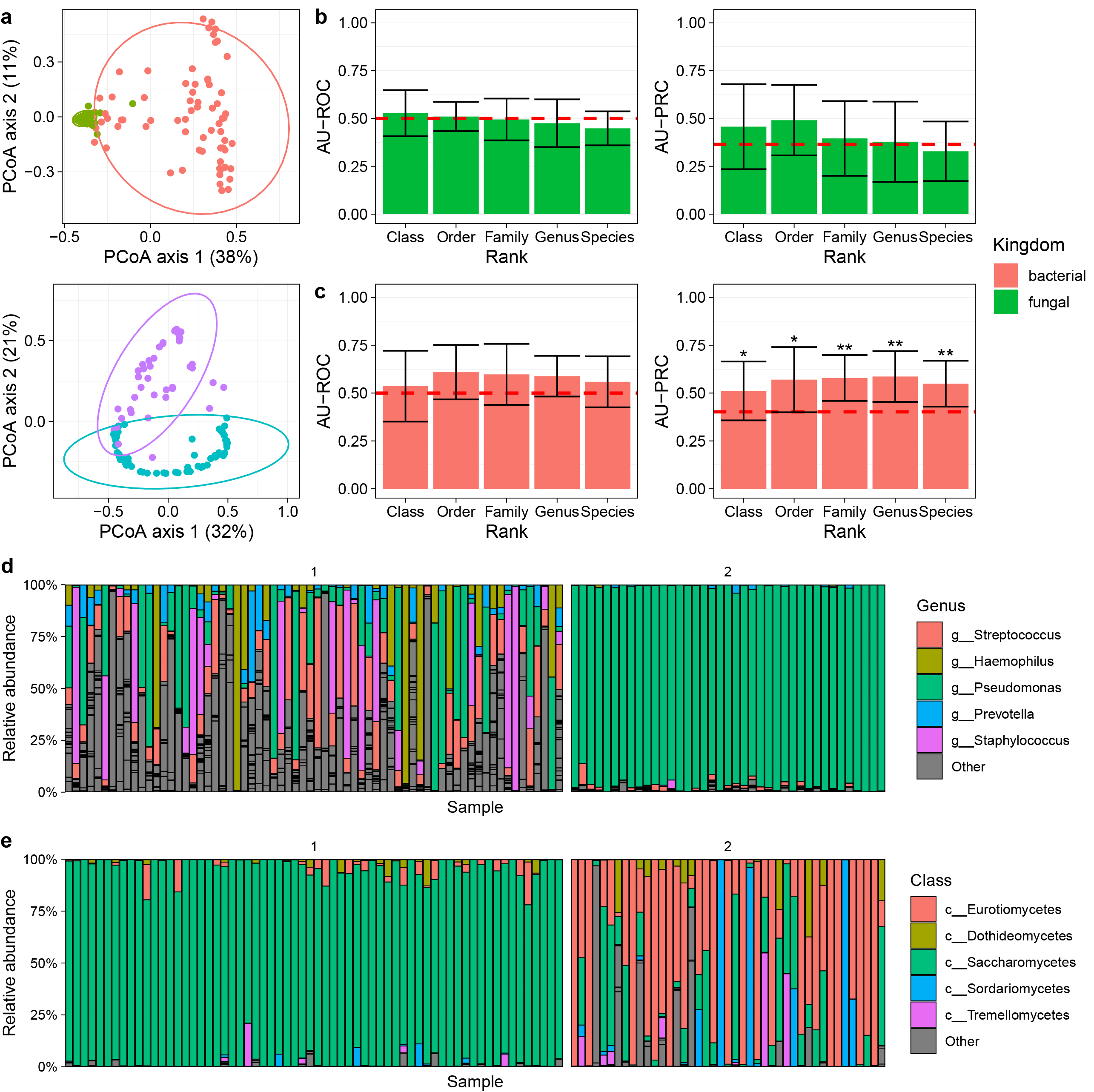
(a, left) Clustering of samples based on bacterial genera abundances identifies two clusters that are separable in Bray-Curtis principle co-ordinate analysis (PCoA) space. (a, right) Clustering using fungal class abundance also finds two clusters that are separable in Bray-Curtis space. The two sets of cluster labels do not correspond to one another nor to clinical labels (see Table 3). (b): Random forest two-sample testing shows that the bacterial cluster assignments are independent of fungal community composition. (c) The fungal cluster assignments show a weak association with bacterial community composition, with only the PR-curves suggesting an association. (d) The bacterial composition of the samples when ordered by cluster clearly shows that they correspond to presence or absence of domination by *Pseudomonas* species. (e) The two fungal clusters are defined by presence or absence of *Saccharomycetes* domination. *: *P* < 0.10, **: *P* < 0.05 for 100 replicates of a label permutation test. *P*-values adjusted using false discovery rate.

Neither the bacterial nor the fungal clusters agree (Adjusted Rand index=-0.01). There is also very low similarity between the cluster labels and clinical labels (Table 3). The ARI values are close to zero, other than for fungal Class cluster and fungal disease status within the CF group (ARI=0.26), however, values still show low levels of agreement. The random forest two-sample testing procedure showed that fungal class is independent of the bacterial community at all levels of agglomeration (Figure 4b). A weak (ROC curves not significantly better than random) association between *Saccharomycetes* domination and bacterial Species, Genus and Family abundance was, however, observed (Figure 4c).

**Table 3.**
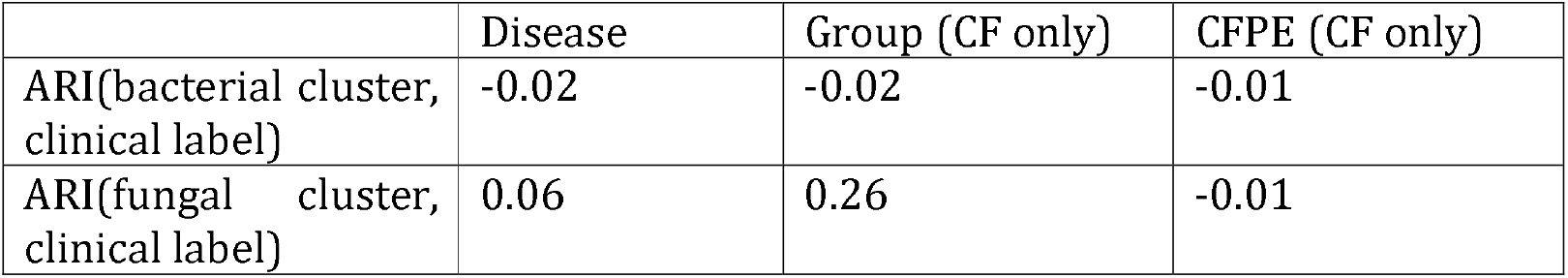
Adjusted Rand Index (ARI). ARI between bacterial/fungal cluster assignments and clinical labels show that there is low similarity between the either set of cluster labels and other clinical labels used in this study. The ARI between the fungal and bacterial clustering labels was −0.01.

## Discussion

Identifying factors predicting the prevalence and severity of chronic respiratory infections may be crucial for improving clinical outcomes in CSLDs. To date the majority of research has been focused on bacterial pathogens. An increasing number of recent studies however, are showing that fungal infection plays a key role in chronic disease progression both independently of and in concert with the bacterial airway community (8, 27, 28). As such, understanding the inter-kingdom association present within the lungs is an essential step towards effective antimicrobial treatments.

A primary motivation of this study was to explore inter-kingdom interactions. Such interactions have been reported previously in both CF and BX (8, 28), as well as in many other settings (29–31). Despite using a range of statistical approaches, we did not however in this present study find strong evidence of such interactions in our dataset (either in general or in relation to CF or BX). Including both kingdoms in the random forest models did not increase the discriminative power of any of the random forest models, while fungal disease status of the CF group was independent of bacterial community composition. Both the correlation and MOFA analysis failed to find evidence of cross-kingdom interactions and instead identified sets of kingdom-specific features that were largely independent of one another. Finally, the sample-wise clustering found that a characteristic feature of fungal community composition (domination by *Saccharomycetes*) was distinct from *Pseudomonas* domination.

Taken together, these results suggest no important cross-kingdom interactions present in this dataset. This is surprising given that both the fungal and bacterial communities are sharing the same niche and so must compete for resources, as well as being affected by common environmental changes. Given the results of previous studies, it is unclear whether such interactions biologically exist but are simply not detectable in this dataset.

A recent study carried out by Hughes *et al*. established that using culture-based methodologies, the known CF pathogens, *Pseudomonas aeruginosa* and *Aspergillus fumigatus* are rarely cultured from the same sample (32). Despite this, our culture independent techniques clearly show the common presence of both *Pseudomonas aeruginosa* and *Aspergillus fumigatus* reads in the samples. It is possible that microbial interactions within the lungs may not be detected by DNA-based methods. Future work may require functional analyses to explore relative microbial gene expression within the lung.

Despite many similarities in the symptoms and treatments of CF and BX, we identified fundamental differences in their microbial communities. Using random forest modelling we found that CF/BX status depends on both fungal and bacterial community composition in this cohort. Furthermore, we found that both communities are equally discriminative of CF/BX status, but the inclusion of both communities in the models does not increase predictive power. This is further evidence that the fungal and bacterial communities are independently distinct between CF and BX and does not provide any evidence of clinically relevant cross-kingdom interactions.

These observed differences between the CF and BX groups are likely to be driven by the physiological differences underlying the individual diseases and their effect on the host environment (33). These differences may also be influenced by age, which is a perfect confounder for disease status in this cohort. It is not possible to correct for this confounding effect as CF is a disease affecting individuals from childhood and BX affects older age groups.

Variable importance analysis with these random forest models identified a set of genera from both kingdoms that are associated with increased likelihood of CF (*Pseudomonas* and *Scedosporium*) and BX (*Penicillium, Neisseria, Campylobacter, Trichocomaceae, Malassezia, Enterobacteriaceae* and *Talaromyces*).

These results are consistent with the known role of *Pseudomonas aeruginosa* as one of the most common pathogens associated with CF lung disease. In non-CF bronchiectasis, *Pseudomonas* infection may be associated with more severe disease (34) but it was not a prominent factor in our BX patients.

Members of the *Neisseria genus* are commonly isolated in the upper respiratory tract with some species being known pathogens (35). Our results may suggest that a pathogenic role for *Neisseria spp.* could be considered for BX and warrants further investigation. Fungal species associated with BX were primarily part of the *Penicillium* genus. Symptomatic infections with *Penicillium* spp. are rare (36) and Penicillum spp. are widely present in the air making it a logical part of the normal respiratory flora.

The microbiota between patients with and without a clinical fungal infection using the random forest pipeline found that the fungal disease status of the CF group was independent of bacterial community composition, but not fungal community composition. The analysis identified several drivers of fungal bronchitis (*Trichocomaceae, Scedosporium, Exophiala* and *Aspergillus*) while also finding that increasing *Candida* decreases the likelihood of a fungal bronchitis diagnosis, consistent with previous findings (7).

Despite the association with the NAFD group, members of the *Candida* genus (including *Candida* albicans and *Candida parapsilosis*) are well-known opportunistic human pathogens, particularly in immunocompromised patients (6). In adult CF patients, *Candida spp*. colonization has been shown to be associated with use of inhaled steroids, diabetes mellitus and antibiotic treatment. Despite these observations the virulence potential of *C. albicans* in CF is still being explored (6). In the current study *Candida spp*. were present with a lower relative abundance in the FB group suggesting that dominance of filamentous fungi may out-compete *Candida spp*. in these patients. More work is therefore needed to understand the role members of the *Candida* genus play in CF disease progression.

Pulmonary exacerbations are major clinical events in patients with CF resulting in lung function decline and clinical disease progression (37). The presence of bacteria and viruses is commonly associated with poor outcomes during CFPE but defining their exact role is challenging. Recent evidence has suggested fungal infections are also associated with increased CFPE although to date few studies have explored this area. A recent publication by Soret *et al*. investigated CFPE using an adapted penalised linear model and cross-sectional data and identified two fungal genera, *Aspergillus* and *Malassezia*, associated with CFPE (8). Our analyses however found that CFPE status was independent of both bacterial and fungal community composition.

The importance of viruses has been shown by the sharp reduction in the incidence of CFPE during the COVID pandemic (38). Future studies should include assays for respiratory viruses, and longitudinal measurements may be used to test if intra-patient variation within the bacterial and fungal communities is a contributing factor.

We further explored the bacterial and fungal communities and their cross-kingdom dependencies using a series of unsupervised statistical analyses. Correlation network analysis identified two blocks of positively co-correlated genera, where each block contained taxa from a single kingdom. Positive correlations are often interpreted to imply mutualistic relationships between organisms and are often observed between phylogenetically related microbes (39). Negative correlations may imply competition within a niche due to competition for resources. These correlations have previously been observed in multi-omic analyses of CFPE (8). Both positive and negative correlations however are often due to unmeasured factors affecting the host environment and so do not necessarily imply a direct relationship between taxa.

Multi-omics analysis using MOFA also found no evidence of cross-kingdom interactions, as the analysis identified a set of kingdom-specific latent factors. The lack of strong cross-kingdom correlation patterns and the results of the MOFA analysis indicates a surprising degree of independence between the two kingdoms although this is inconsistent with previous studies that have indicated a number of inter-kingdom interactions existing within the lung (8, 28). Unsupervised sample-wise clustering analysis identified characteristic features of the dataset identifying two bacterial clusters at the Genus level and two fungal clusters at the Class level. These two sets of cluster labels had low similarity with one another and with the clinical labels from the random forest analyses, suggesting that the relevant structure of the communities may be primarily due to other (possibly environmental) factors. The bacterial clusters were driven by dominance of *Pseudomonas* within individual samples. *Pseudomonas* was also identified by the random forest analysis as being associated with CF, but these clustering results suggest that Pseudomonas-dominance is not the only predictor of CF in this cohort. The bacterial cluster label was independent from fungal community composition, providing additional evidence of independence between the bacterial and fungal communities.

Inter-kingdom correlations were generally weaker than those within either kingdom (measured by proportion of significant correlations at different significance thresholds). This further indicates that intra-kingdom interactions may play a minor role in these subjects. In addition, correlation patterns between bacterial genera were stronger than those between fungal genera.

Our analysis has several limitations that should be considered when interpreting the results. Machine learning is a powerful tool for exploring microbial interactions and drivers of disease, but understanding the limitations of the models is vital for interpretation. Most importantly, associations identified by machine learning models such as random forest do not imply causal links. Furthermore, the importance scores from random forests should be interpreted with care. Using multiple random forest variable importance scores and transformations in the differential abundance analysis reduces the danger of spurious associations but does not provide a framework that allows quantitative statements to be made.

A further limitation of this study is the use of 16S rRNA gene sequencing and ITS2 sequencing for the exploration of these communities. This technology allows us to understand the microbial community present within the lung but provides no information on their activity or function.

## Conclusions

Our study suggests that the role the fungal microbiota play in chronic respiratory disease is independent of that played by the bacterial microbiota. Longitudinal studies are required to understand the full impact of fungal infection in CF and BX. Importantly improvements in clinical diagnosis of fungal infections, whether by sequence analysis, transcriptomics, or advanced cultures, could underpin the improvement of patient outcomes. While further work is required to fully understand microbial interaction within the lung, our data suggests that inter-kingdom interactions may not be a major driver of patient outcomes particularly those associated with fungal infection.

## Declarations

### Availability of data and material

Sequencing data is freely assessable through the European nucleotide database, bacterial data can be accessed under project number PRJEB33064, while the fungal data can be accessed under project number PRJEB33434.

Data analysis scripts are freely available on figshare, under the project “Machine learning for exploring microbial inter-kingdom associations in Cystic Fibrosis and Bronchiectasis” DOIs: https://doi.org/10.6084/m9.figshare.17897708 and https://doi.org/10.6084/m9.figshare.17942543.

### Authors’ contributions

All authors listed provided substantial contributions to this manuscript. Study concept and design; MFM, WOCC, SF, LC, NJS, MRL, MJC. Acquisition of data; IF, PJ, LC, MJC. Statistical analysis and interpretation of the data; LC, JIH. Drafting of the manuscript; LC, JIH, WOCC, SF. Critical revision of the manuscript was carried out by all authors.

