## Supplementary information for "Cross-kingdom analysis of microbial communities in Cystic Fibrosis and Bronchiectasis"

**Supplementary Methods**

**Decontamination**

After initial sample processing, in QIIME1.9, data was moved to R for all further analyses. Pre-processing and decontamination were carried out separately for bacterial and fungal data. Mock communities and negative controls were examined prior to decontamination.

*Bacterial decontamination*

Decontam version 1.1.2 was used to identify contaminating OTUs. All decontam methods were investigated. Bacterial mock communities were comprised of 35 known bacterial species, including known respiratory organisms and *Vibro natriegens.*  *V. natriegens* was included as a control to indicate contamination from the mock community and/or barcode switching in the sequencing. Vibro OTUs were indicated as contamination using the frequency method in decontam suggesting this was the most appropriate method for the identification of contaminants in this dataset. All OTUs from the Vibrio genus were removed before final application of decontam to the dataset and the removal of 56 contaminant OTUs.

Finally, OTUs unassigned at the kingdom level, mitochondria and the known contaminants from Rhodobacterales, Rhizobiales, Oxalobacteraceae and Cyanobacteria were removed from the dataset. These OTUs were in low abundance and did not affect the results obtained from further analysis. Finally, low abundant OTUs were removed by removing OTUs with less than 20 reads.

*Fungal decontamination*

As previously described in Cuthbertson *et al*. 2020 (1), the decontam frequency method was found to be the most appropriate to identify fungal contaminants. Seventy-four OTUs were identified and removed from further analysis. Indicator species analysis, using the package indicspecies version 1.7.6, was carried out to identify OTUs associated with Extraction kit and controls.

As with the bacterial mock, the fungal mock community included a control organism *Batrachochytrium dendrobatidis*. Batrachochytrium OTUs were removed from further analysis. Finally, low abundant OTUs with less than 20 reads were removed from further analysis.

**Results**

**Ecological analysis**

*Differences between diseases*

An ecological analysis of the rarefied OTU counts (Fungi; n = 2,542, Bacteria; n = 2,357) was also performed. Both bacterial and fungal diversity were significantly higher in patients with BX than CF (Bacteria; richness, W = 1,507, effect size = 0.369, *P* < 0.001, Shannon, W = 1,545, effect size = 0.396, *P* < 0.001, Simpson, W = 1,549, effect size = 0.399, *P* < 0.001. Fungi; richness, W = 1,603, effect size = 0.438, *P* < 0.001, Shannon, W = 1,522, effect size = 0.379, *P* < 0.001, Simpson, W = 1,465, effect size = 0.338, *P* < 0.001). No significant difference in dominance, however, was observed in either bacterial or fungal communities between diseases (Bacteria; W = 786, effect size = 0.151, *P* = 0.118. Fungi; W = 781, effect size = 0.155, *P* = 0.108). Although bacterial biomass was significantly lower in the CF group compared to BX (W = 1,100, effect size = 0.241, *P* = 0.018), no significant difference in fungal biomass was observed (W = 706, effect size = 0.112, *P* = 0.126). Differences in both bacterial and fungal community composition, calculated by PERMANOVA, showed significant differences between CF and BX. The effect sizes however were extremely small (bacterial, R^2^ = 0.066, *P* < 0.001; fungal R^2^ = 0.028, *P* = 0.004).

*Differences between clinical fungal disease groups*

Participants within the study cohort were classified into four clinically defined fungal disease groups (see Main Paper Table 2): (i) fungal bronchitis (FB), (ii) Allergic bronchopulmonary aspergillosis (ABPA), (iii) chronic necrotising pulmonary aspergillosis (CNPA, BX only) and (iv) non-tuberculosis mycobacteria (NTM). NTM patients were included as a subgroup due to the strong link between NTM infection and fungal disease. [7 38, 39]. A control group of patients presenting with no active fungal disease (NAFD) were also included.. Due to the small number of patients with BX in the study (n = 24) the analysis was confined to the CF (n=83) patient samples. Within the CF group, we compared the two largest fungal disease groups, which were FB (n=20) and NAFD (n=39).

We observed significant differences in fungal alpha diversity between the 39 CF patients with NAFD and the 20 with FB (Biomass: W = 522, effect size = 0.275, *P* = 0.03, richness; W = 211, effect size = 0.374, *P* < 0.004; Shannon: W = 204, effect size = 0.387, *P* = 0.002; Simpson: W = 213, effect size = 0.368, *P* = 0.004). We found no significant differences in bacterial biomass or diversity between fungal disease groups.

Fungal beta diversity using Bray-Curtis similarity was also significantly different between the CF and BX groups (PERMANOVA R^2^ = 0.077, *P* = 0.004), although the R^2^ effect size was small.

*CFPE*

Thirty-six of the 83 CF patients in the study were defined as experiencing CFPE at the time of sampling. There was no significant difference in bacterial or fungal biomass or alpha diversity measures (Richness, Shannon, Simpson) (*P* > 0.1) between CFPE subjects and those that were stable using Wilcoxon rank sum test.

Tables and figures

**Table S1**: Uploaded separately, supplementary_table_1.xlsx

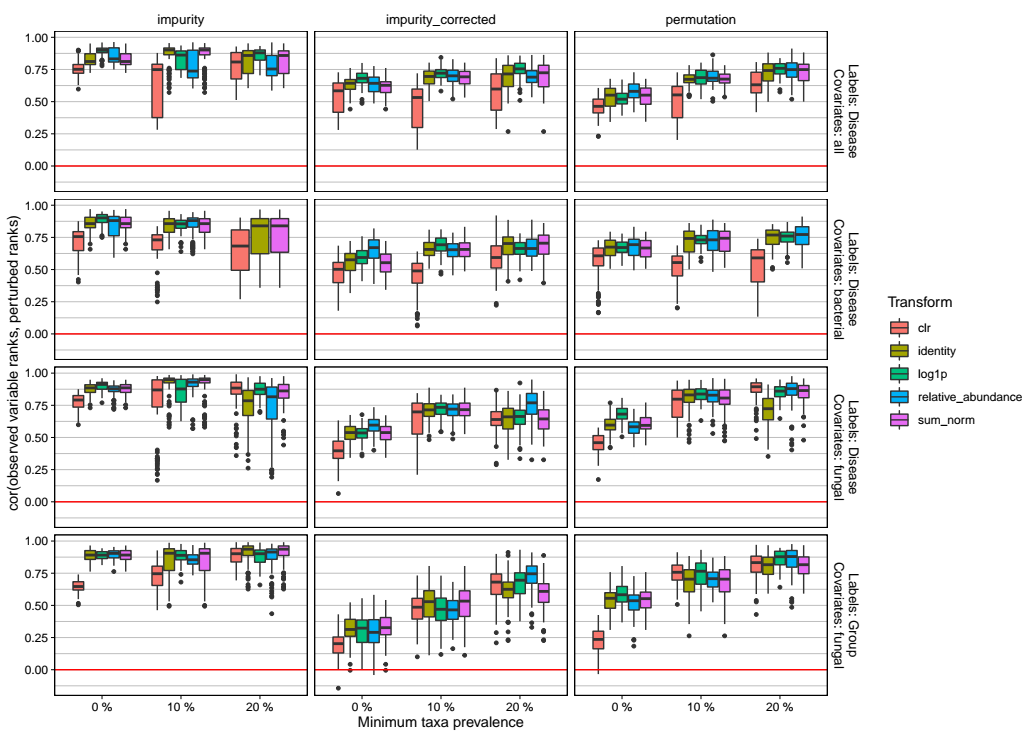

**Figure S1**: Spearman correlation between variable rankings for random forests trained using the observed dataset and 100 perturbed datasets (removal of 10% of samples). Removing rare taxa increases the stability of the variable importance rankings under these perturbations.

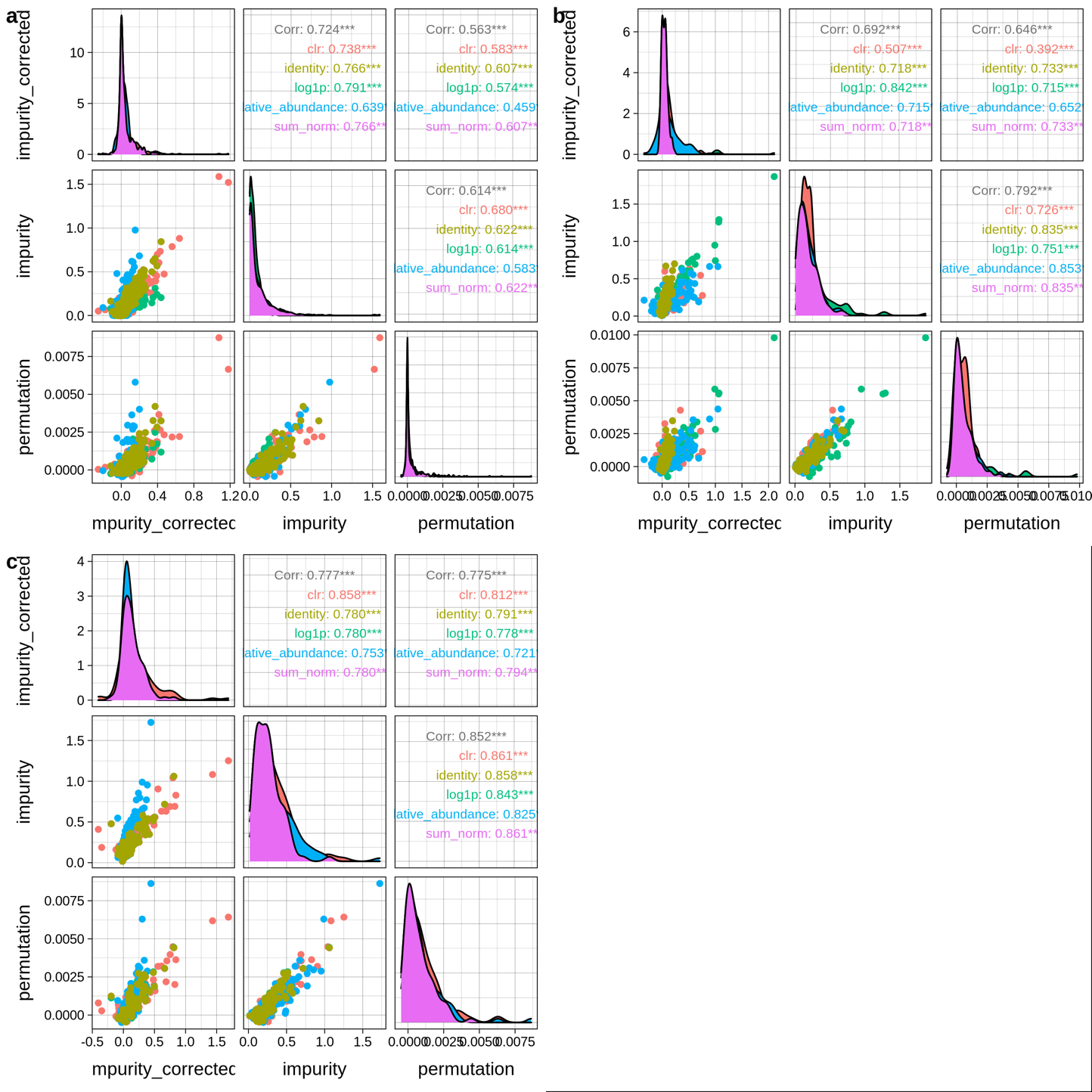

**Figure S2**: Agreement between the random forest variable importance methods under different transformations. Model: predicting disease status from bacterial and fungal genera. (a): no taxa removed, (b): taxa present in fewer than 10% of samples removed, (c): taxa present in fewer than 20% of samples removed.

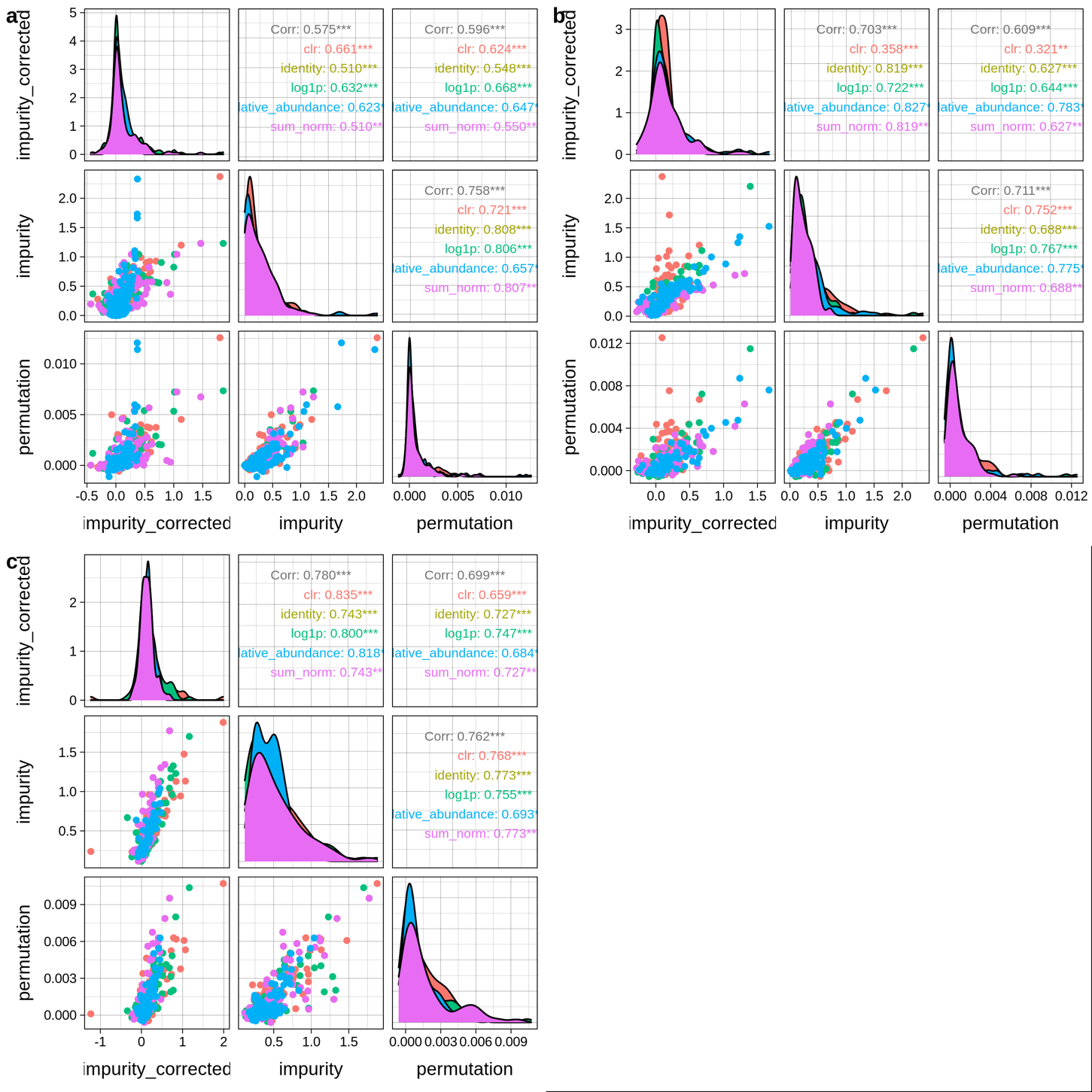

**Figure S3**: Agreement between the random forest variable importance methods under different transformations. Model: predicting disease status from bacterial genera. (a): no taxa removed, (b): taxa present in fewer than 10% of samples removed, (c): taxa present in fewer than 20% of samples removed.

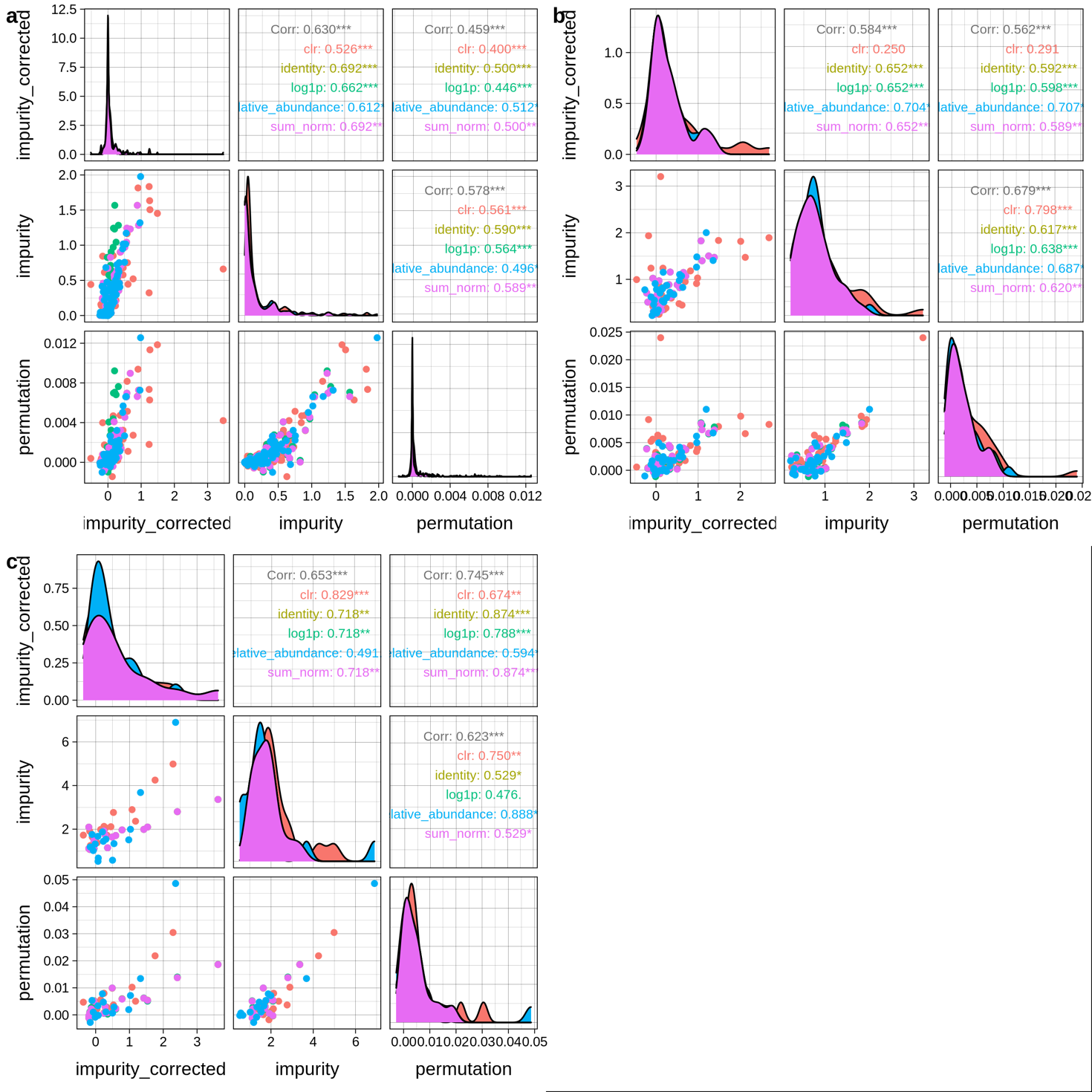

**Figure S4**: Agreement between the random forest variable importance methods under different transformations. Model: predicting disease status from fungal genera. (a): no taxa removed, (b): taxa present in fewer than 10% of samples removed, (c): taxa present in fewer than 20% of samples removed.

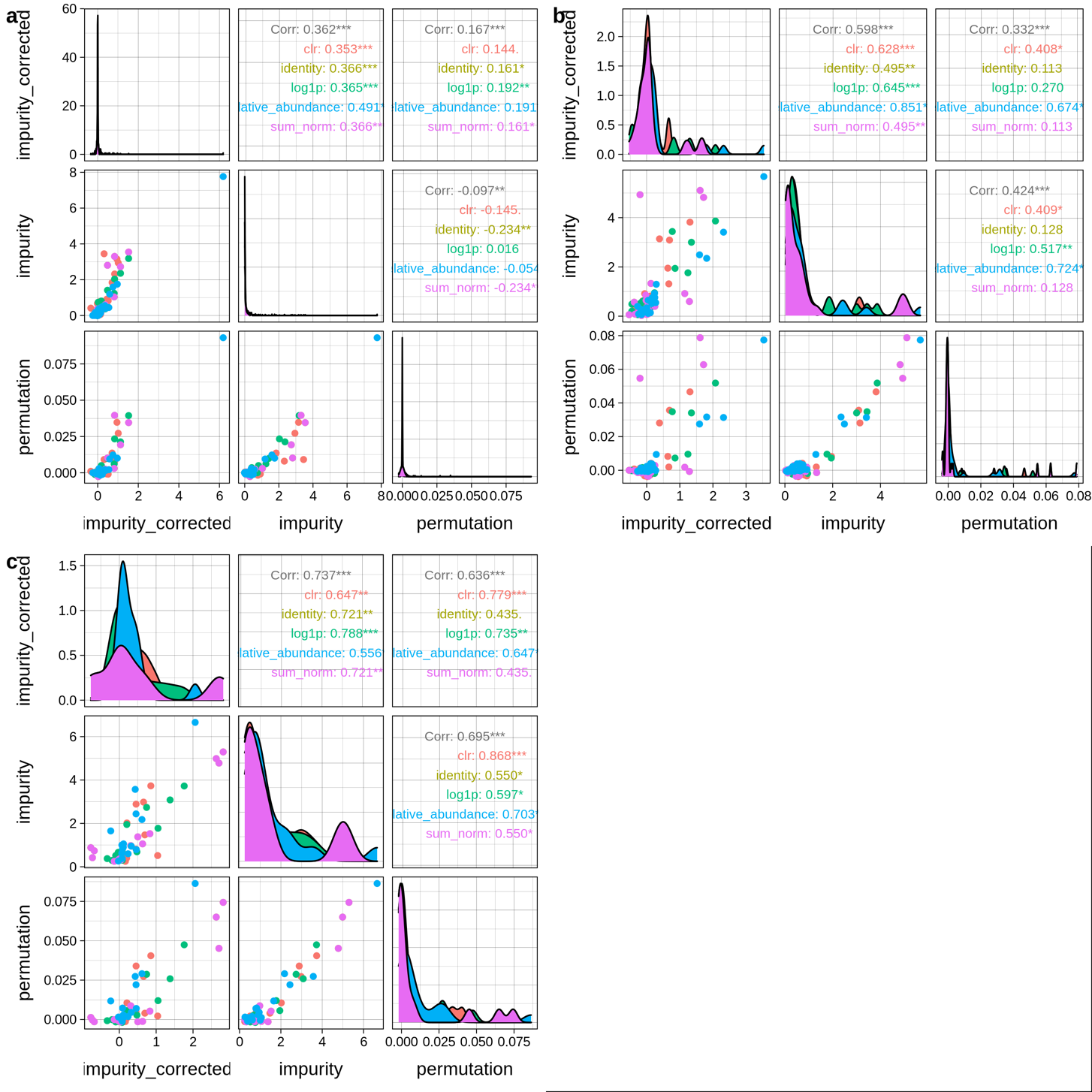

**Figure S5**: agreement between the random forest variable importance methods under different transformations. Model: predicting fungal disease status (CF group only) from fungal genera. (a): no taxa removed, (b): taxa present in fewer than 10% of samples removed, (c): taxa present in fewer than 20% of samples removed.

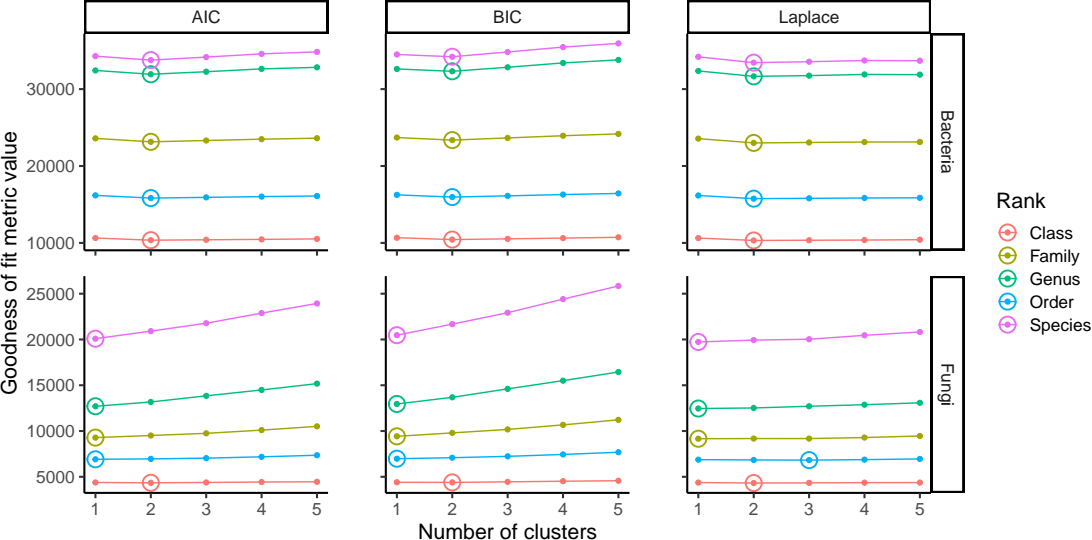

**Figure S6**: Goodness-of-fit for (sample-wise) Dirichlet Multinomial clustering of the bacterial (top row) and fungal (bottom row) abundances at different levels of agglomeration. Each column contains a different goodness-of-fit metric (AIC: Akaike information criterion, BIC: Bayesian information criterion) where lower values indicate a better fit. The number of clusters that minimises the metric is circled. These results indicate that the 107 samples form two clusters based on their bacterial community composition and one cluster based on their fungal community composition.
